# Regulation of the hypertonic stress response by the 3’ mRNA cleavage and polyadenylation complex

**DOI:** 10.1101/2023.01.23.525244

**Authors:** Sarel J. Urso, Anson Sathaseevan, W. Brent Derry, Todd Lamitina

**Author notes:** Address correspondence to Todd Lamitina, Ph.D., Children’s Hospital of Pittsburgh of the University of Pittsburgh Medical Center, Departments of Pediatrics and Cell Biology, 4401 Penn Avenue, Rangos 7122, Pittsburgh, PA 15224.

## Abstract

Maintenance of osmotic homeostasis is one of the most aggressively defended homeostatic setpoints in physiology. One major mechanism of osmotic homeostasis involves the upregulation of proteins that catalyze the accumulation of solutes called organic osmolytes. To better understand how osmolyte accumulation proteins are regulated, we conducted forward genetic screen in *C. elegans* for mutants with no induction of osmolyte biosynthesis gene expression (Nio mutants). *nio-3* mutants encoded a missense mutation in *cpf-2*/CstF64 while *nio-7* mutants encoded a missense mutation in *symk-1*/Symplekin. Both *cpf-2* and *symk-1* are nuclear components of the highly conserved 3’ mRNA cleavage and polyadenylation complex. *cpf-2* and *symk-1* block the hypertonic induction of *gpdh-1* and other osmotically induced mRNAs, suggesting they act at the transcriptional level. We generated a functional auxin-inducible degron (AID) allele for *symk-1* and found that acute, post-developmental degradation in the intestine and hypodermis was sufficient to cause the Nio phenotype. *symk-1* and *cpf-2* exhibit genetic interactions that strongly suggest they function through alterations in 3’ mRNA cleavage and/or alternative polyadenylation. Consistent with this hypothesis, we find that inhibition of several other components of the mRNA cleavage complex also cause a Nio phenotype. *cpf-2* and *symk-1* specifically affect the osmotic stress response since heat shock-induced upregulation of a *hsp-16*.*2::GFP* reporter is normal in these mutants. Our data suggest a model in which alternative polyadenylation of one or more mRNAs is essential to regulate the hypertonic stress response.

## Introduction

mRNA processing plays a key role in the response to diverse types of environmental stress. For example, post-transcriptional splicing of the *xbp-1* mRNA by activated IRE1 endonuclease is the key step in activation of the endoplasmic reticulum (ER) stress response (CALFON *et al*. 2002). In the DNA damage response, localized mRNA splicing plays a key role in the active repair of double strand breaks (SHANBHAG *et al*. 2010; PEDERIVA *et al*. 2016). Recently, 3’ mRNA cleavage components from transcriptionally active genomic loci were found to undergo hyperosmotic stress induced phase separation, which was associated with a global impairment of transcriptional termination (JALIHAL *et al*. 2020). Whether or not 3’ mRNA cleavage or other types of mRNA processing play a more direct functional role in the regulation of the hyperosmotic stress response is unknown.

In eukaryotes, the 3’ ends of ∼30-60% of all mRNAs are cleaved and polyadenylated at more than one polyadenylation site (PAS), which results in mRNAs with different 3’ ends (STEBER *et al*. 2019; YUAN *et al*. 2021). Much like alternative splicing, alternative polydenylation (APA) also leads to the diversification of mRNA transcripts through two distinct mechanisms. First, 3’ UTR APA gives rise to mRNAs with different length 3’ UTRs, which can provide isoform specific binding sites for mRNA regulators such as micro RNAs (miRNAs) and RNA binding proteins (MAYR AND BARTEL 2009). Second, upstream region APA (UR-APA) can include alternative exons that, when translated, give rise to proteins with unique sequences and/or domains (ALT *et al*. 1980). The 3’ mRNA end processing complex is composed of ∼85 proteins that regulate mRNA cleavage, polyadenylation, and APA (SHI *et al*. 2009). Biochemical and structural features of the core complex of ∼20 protein are relatively well described (PANCEVAC *et al*. 2010; CLERICI *et al*. 2017; CLERICI *et al*. 2018) and new sequencing and bioinformatics approaches are enabling the identification of mRNA targets of APA (HA *et al*. 2018; WANG *et al*. 2018). While APA is associated with aspects of cellular physiology, such as cellular stress responses (CHANG *et al*. 2018; ZHENG *et al*. 2018), the physiological roles of 3’ mRNA cleavage/APA are poorly understood, largely because genetic null alleles of most 3’ mRNA cleavage components are lethal, although some important exceptions do exist (SUBRAMANIAN *et al*. 2021).

In *C. elegans*, hypertonic stress activates a gene expression program that restores cell volume and protects against hypertonicity-induced macromolecular damage (ROHLFING *et al*. 2010). Like all cells and organisms, *C. elegans* uses solutes called organic osmolytes to restore volume and provide protection (LAMITINA *et al*. 2004; LAMITINA *et al*. 2006; BURKEWITZ *et al*. 2012). The major *C. elegans* osmolyte is glycerol, which is rapidly accumulated following exposure to hypertonicity, in part through the upregulation of the glycerol-3-phosphate dehydrogenase *gpdh-1* (LAMITINA *et al*. 2004). Genetic screens for regulators of *gpdh-1* expression have identified many positive and negative regulators of this stress response pathway (LAMITINA *et al*. 2006; WHEELER AND THOMAS 2006; ROHLFING *et al*. 2011; DODD *et al*. 2018; URSO *et al*. 2020; WIMBERLY AND CHOE 2022). However, it is not known whether components of the 3’ mRNA cleavage complex play a functional role in the regulation of the hypertonic stress response.

Using unbiased forward genetic screening for mutants that exhibit no induction of *gpdh-1* expression in response to hypertonicity, we identified viable hypomorphic alleles of two 3’ mRNA cleavage complex genes, *cpf-2* and *symk-1. cpf-2* and *symk-1* physiologically regulate the transcriptional induction of multiple hypertonicity induced mRNAs through their activity in the hypodermis and intestine. CPF-2 and SYMK-1 proteins co-localize within the nucleus of all cells and form subnuclear puncta in response to hypertonicity. Both *symk-1* and *cpf-2* mutants do not significantly alter general mRNA polyadenylation but instead exhibit phenotypes consistent with inhibition of APA. Consistent with this hypothesis, inhibition of several other 3’ mRNA cleavage complex genes also blocks hypertonic induction of *gpdh-1* transcription. *cpf-2* and *symk-1* are not required for the transcriptional induction of a heat shock inducible GFP reporter, suggesting that their roles in regulating stress response pathways may be specific. Our findings reveal a physiological role for the 3’ mRNA cleavage complex complex in the hypertonic stress response and provide new genetic tools that should facilitate the *in vivo* study of this essential mRNA regulatory mechanism in developmental and tissue-specific contexts.

## Results

In a forward genetic screen for mutants with *<u>n</u>*o *i*nduction of the *<u>o</u>*smolyte biosynthetic transcriptional reporter *gpdh-1p::gfp* reporter (Nio muants), we identified *nio-3(dr16)* and *nio-3(dr23)*. Mapping, complementation testing, and whole genome sequencing revealed that *dr16* caused a G42E missense mutation in the *cpf-2* gene while *dr23* caused a R580W missense mutation in the *symk-1* gene (Fig. 1A). Both alleles were genetically recessive and segregated Nio animals from heterozygous hermaphrodites close to expected single gene Mendelian ratios (*dr16* – 24/111, 21.6%; *dr23* – 22/145, 15.2%). *cpf-2* and *symk-1* encode two proteins that are core components of the 3’ mRNA cleavage and polyadenylation complex and are known to interact in mammalian cells (RUEPP *et al*. 2011). This complex recognizes polyadenylation signals in the mRNA and directs mRNA cleavage and transcriptional termination through a multi-subunit protein complex (SHI *et al*. 2009). Several additional experiments show that these non-null mutations in *cpf-2* and *symk-1* are the basis for the Nio phenotype. First, post-embryonic RNAi of both *cpf-2* and *symk-1* also caused a Nio phenotype (Fig S1).

**Figure 1.**
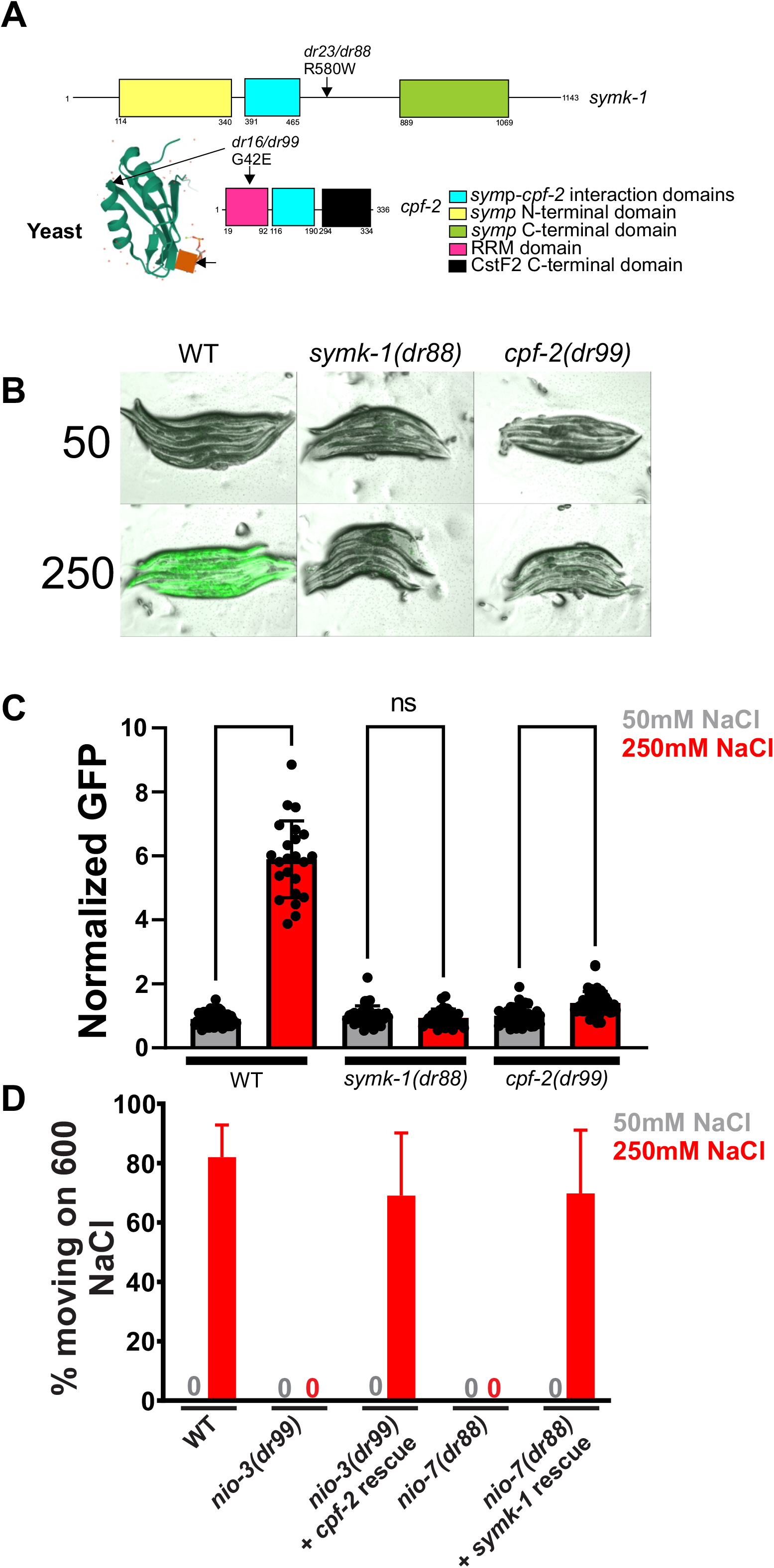
*nio-3* and *nio-7* are caused by missense mutations in the polyadenylation complex genes *cpf-2* and *symk-1*. A) Protein domains found in SYMK-1 and CPF-2. The location of each missense mutation is indicated. For *cpf-2*, we used the available crystal structure of the APA complex to map the location of the conserved G42 residue on the yeast homolog of *cpf-2*, Rna15 (PANCEVAC *et al*. 2010). B) Overlayed transmitted light and GFP images of CRISPR-engineered animals carrying missense Nio alleles of either *symk-1* or *cpf-2* exposed to either 50 or 250 mM NaCl for 18 hours. C) COPAS quantification of animals imaged in B. Bar graphs show the mean ± S.D., with individual data points overlayed. D) Hypertonic adaptation of CRISPR Nio alleles with or without a genomic DNA rescue transgene. Data are the mean ± S.D. of 4-5 independent replicates of 10-20 worms per replicate.

Second, extrachromosomal arrays carrying either fosmids or genomic DNA PCR products encompassing either the *cpf-2* and *symk-1* loci rescue *dr16* and *dr23* accordingly (Fig. S2). Third, CRISPR engineered alleles of *dr16* and *dr23* that recreated each missense mutation in a non-mutagenized *drIs4* background also blocked hypertonic induction of *gpdh-1p::GFP* (Fig. 1B,C) and eliminated the ability to adapt to normally lethal hypertonic environments (Fig. 1D). Notably, null or strong loss of function alleles of both *cpf-2* and *symk-1* are embryonic/larval lethal (STEBER *et al*. 2019) whereas the Nio alleles are viable and fertile under isotonic conditions. Taken together, these data show that *nio-3(dr16)* and *nio-7(dr23)* are caused by non-null hypomorphic alleles in *cpf-2* and *symk-1*.

The Nio screen utilized the *drIs4* reporter strain to identify mutants with defective transcriptional responses to hypertonic stress. To determine if these alleles also affected endogenous mRNAs, we performed qPCR on several genes previously shown to be upregulated by hypertonicity (ROHLFING *et al*. 2010). Hypertonic upregulation of endogenous *gpdh-1* mRNA, as well as another hypertonicity upregulated gene *hmit-1*.*1* and the transgene derived *gfp* mRNA, was strongly attenuated in both *cpf-2(dr99)* and *symk-1(dr88)* (Fig. 2). However, upregulation of *nlp-29*, an antimicrobial peptide previously shown to be induced by infection, wounding, and hypertonicity (PUJOL *et al*. 2008), continued to be upregulated to wild type levels in *cpf-2(dr99)* and *symk-1(dr88)*. Therefore *cpf-2* and *symk-1* are required to upregulate some but not all hypertonicity responsive mRNAs.

**Figure 2.**
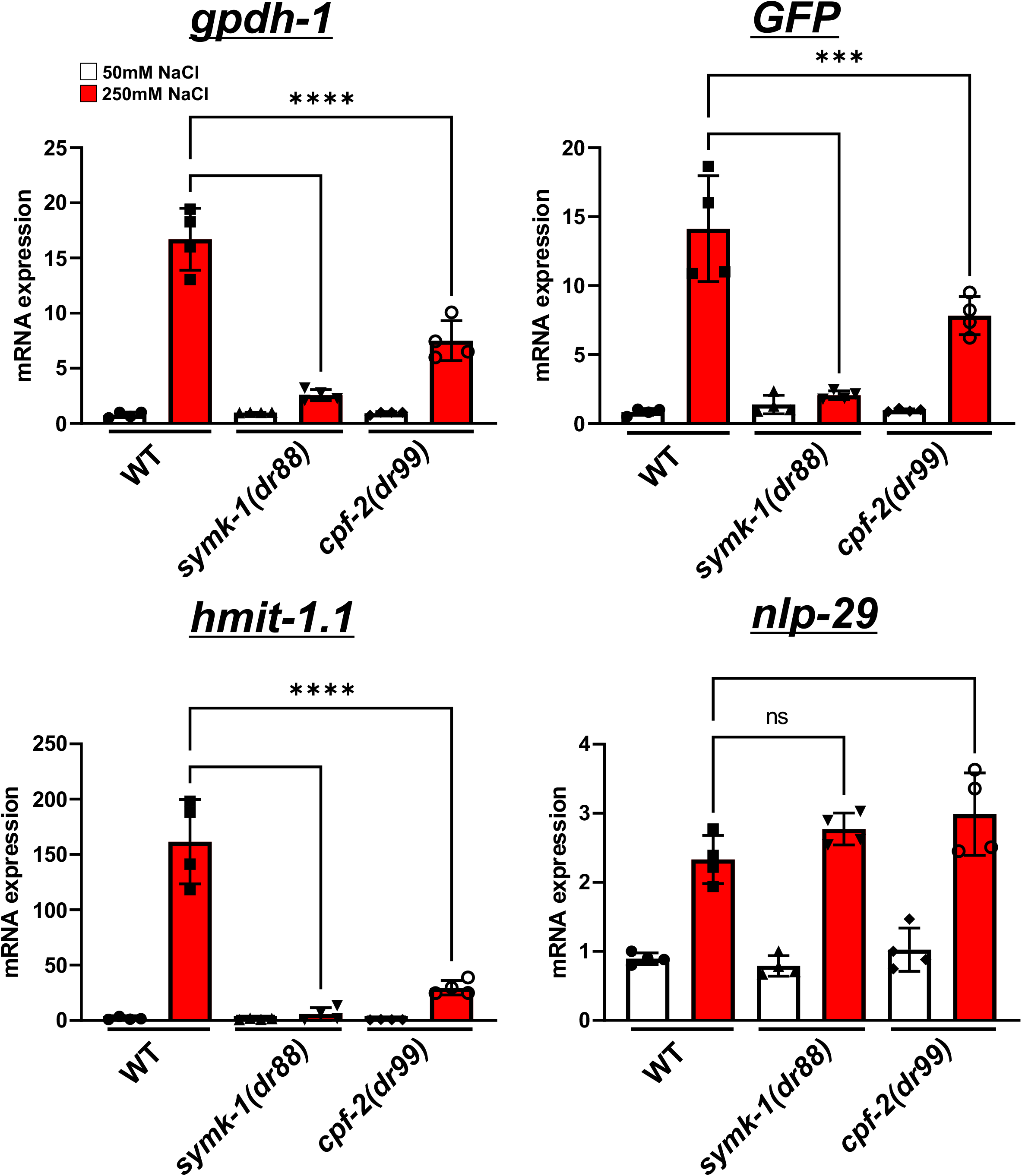
*cpf-2* and *symk-1* are required for hypertonic stress induced mRNA expression. qPCR of the *gpdh-1, gfp* (from the *drIs4* reporter), *hmit-1*.*1*, and *nlp-29* mRNAs in wild type, *symk-1(dr88)*, and *cpf-2(dr99)* young adult animals exposed to 50 mM or 250 mM NaCl NGM. Data are the mean ± S.D. of 4 independent biological replicates. Individual data points are plotted. ****- p<0.0001, ***-p<0.001, *-p<0.05, ns-p>0.05, One-way ANOVA with Sidak multiple comparisons test.

*gpdh-1* is upregulated by hypertonic stress in the intestine and the hypodermis (LAMITINA *et al*. 2004; LAMITINA *et al*. 2006). If *cpf-2* and *symk-1* function cell autonomously, then they should also be required in the intestine and hypodermis to upregulate *gpdh-1* in response to hypertonicity. To test this hypothesis, we generated auxin-inducible degradation (AID*) alleles for both *cpf-2* and *symk-1* using CRISPR/Cas9 methods. The *AID*-GFP-cpf-2* allele was sick and exhibited low fecundity, suggesting the presence of non-specific *cpf-2* degradation and was not studied further. However, the *symk-1::GFP::AID\** allele exhibited qualitatively normal growth and fertility in the absence of auxin, suggesting this allele is functional. We generated homozygous *symk-1::GFP::AID\** alleles in genetic backgrounds expressing the TIR1 E3 ligase in all cells, hypodermis, intestine, or hypodermis and intestine (ASHLEY *et al*. 2021). While *symk-1::GFP::AID\** animals expressing global TIR1 robustly expressed nuclear GFP in all somatic and germline cells in the absence of auxin, exposure of animals to 0.5mM auxin led to loss of the GFP signal in all tissues within 3-4 hours (Fig. S3). Additionally, 100% of the progeny derived from L4 animals exposed to 0.5mM auxin paired with global expression of TIR1 were embryonic or larval lethal, suggesting that auxin-induced depletion of *symk-1* mimics the *symk-1* null phenotype.

Next, we examined induction of the *gpdh-1p::GFP* reporter in each TIR1 strain in the absence of auxin. GFP expression was induced in the intestine and hypodermis to similar levels as wild type (Fig. S4). When animals were exposed to auxin starting as L1s, hypertonic induction of GFP was inhibited in the global TIR1 strain as much as in the *symk-1(dr88)* allele (Fig. 3A,B). Hypodermal depletion of *symk-1* led to an almost complete inhibition of GFP induction by hypertonicity, while intestinal depletion of *symk-1* had a minor effect. However, simultaneous depletion of *symk-1* in both the intestine and hypodermis resulted in a complete block in hypertonicity *induced gpdh-1::GFP* expression. We next utilized the *symk-1 AID\** allele and hypodermal/intestinal TIR1 to examine the temporal requirements for *symk-1* in the activation of *gpdh-1*. Exposure to auxin for four hours in Day 1 adult animals immediately prior to exposure to hypertonicity was sufficient to block *gpdh-1p::GFP* expression (Fig. 3C). Taken together, these data show that *symk-1* functions cell autonomously in the hypodermis and intestine to regulate hypertonic *gpdh-1* induction. Additionally, they show that *symk-1* has a post-developmental physiological role in this process.

**Figure 3.**
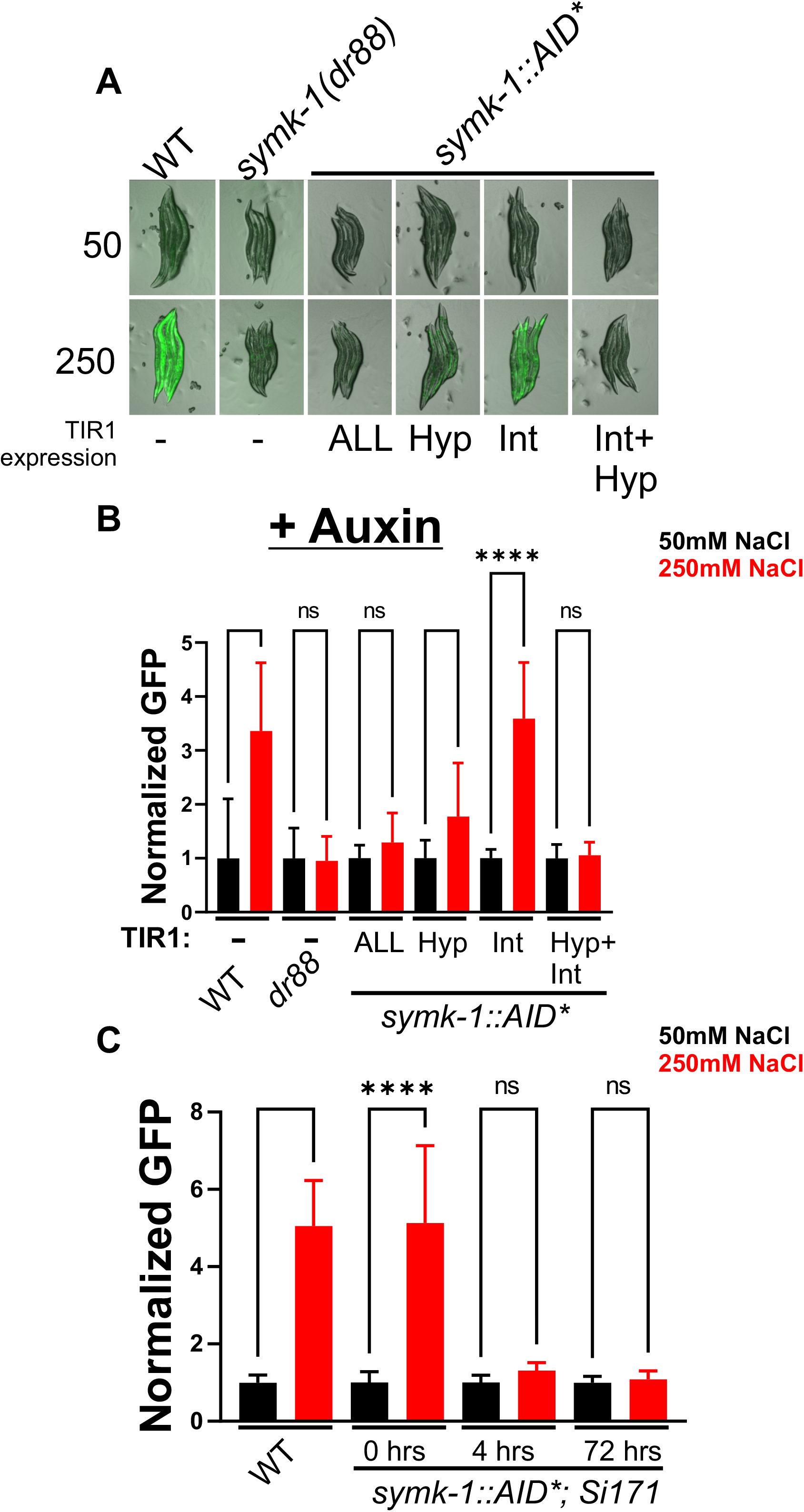
*symk-1* is required post-developmentally in the hypodermis and intestine for the Nio phenotype. A) Overlayed GFP and transmitted light images of *gpdh-1p::gfp* expression following 24 hours of exposure to 50mM or 250mM NaCl. Animals express TIR1 in the indicated tissues and were exposed to 1 mM K-NAA from the L1 stage. B) COPAS Biosort quantification of GFP expression from animals shown in A. Data shown are mean ± S.D. N=41-89 animals per sample. ****-p<0.0001, ns-p>0.05, One-way ANOVA with Dunn’s multiple comparison test. C) COPAS Biosort quantification of animals with TIR1 expression in the hypodermis and intestine (*cpSi171*) exposed to 1 mM K-NAA for 0, 4, or 72 hours prior to exposure to 50mM or 250mM NaCl. Data shown are mean ± S.D. N=35-61 animals per sample. ****-p<0.0001, ns-p>0.05.

If *cpf-2* and *symk-1* interact within the 3’ mRNA cleavage complex in *C. elegans* as they do in mammalian cells (RUEPP *et al*. 2011), we predicted that they should exhibit spatial co-localization. Indeed, GFP and RFP alleles of endogenous *cpf-2* and *symk-1*, respectively, showed that both proteins are expressed ubiquitously, exhibit co-localization in the nucleoplasm, and are excluded from the nucleolus (Fig. 4). Following exposure to hypertonicity, both proteins formed subnuclear puncta that exhibited complete co-localization. In the same screen that isolated *cpf-2* and *symk-1*, we also identified mutations in the O-GlcNAc transferase homolog *ogt-1*, suggesting that these three proteins could be components of the same complex that regulates the hypertonic stress response. However, our data were inconsistent with this model. We found that that the CPF-2 and SYMK-1 puncta were distinct from previously reported OGT-1 puncta induced by hypertonic stress (Fig S5) (URSO *et al*. 2020). Hypertonicity induced CPF-2/SYMK-1 puncta were most apparent in the hypodermis and to a lesser extent in the intestine, while no puncta were observed in germline nuclei (Fig. 4). Introduction of the Nio allele G42E into *TagRFP*-*cpf-2* did not alter CPF-2 protein abundance, nuclear localization, puncta formation, or co-localization with SYMK-1.

**Figure 4.**
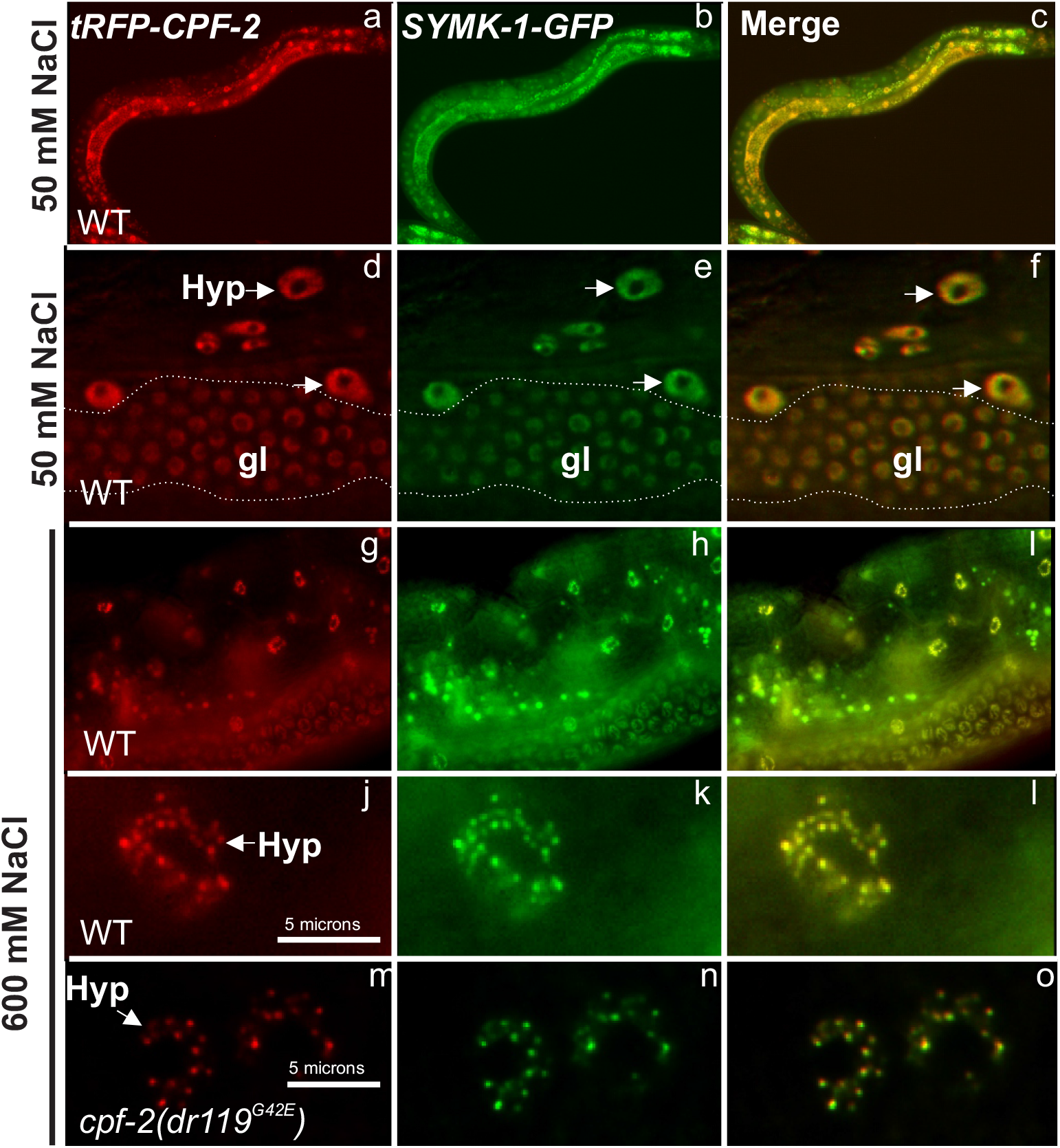
CPF-2 and SYMK-1 proteins colocalize into subnuclear puncta following exposure to hypertonic stress. Expression of CRISPR/Cas9-tagged alleles of *Tag-RFP::cpf-2* (a) *symk-1::gfp* (b), and the merged image (c). d-f – higher resolution images of hypodermal (hyp arrows) and germline (gl) nuclei revealing the non-nucleolar nucleoplasmic concentration of tagged SYMK-1 and CPF-2 proteins. Dotted lines delineate the boundaries of the germline. g-i – Exposure of animals to 600 mM NaCl for 1 hour leads to reorganization of SYMK-1 and CPF-2 into subnuclear foci, primarily in the hypodermis. j-l – higher resolution of hypodermal nuclei with SYMK-1 and CPF-2 foci. m-o-higher resolution of hypodermal nuclei with SYMK-1 and CPF-2 carrying the G42E Nio point mutation.

The 3’ mRNA cleavage and polyadenylation complex is required for recognition of DNA-encoded polyadenylation signal sequences, transcriptional termination via mRNA cleavage, and mRNA polyadenylation. Mutation of *cpf-2* and *symk-1* could cause a Nio phenotype by disrupting these or other processes. Three pieces of data suggest that the Nio alleles do not disrupt global mRNA cleavage and/or polyadenylation at a significant level. First, total RNA levels are not decreased in either *cpf-2(dr99)* or *symk-1(dr88)* (Fig. S6). Second, polyA patterns and lengths for several mRNAs are not altered in either *cpf-2(dr99)* or *symk-1(dr88)* (Fig. S7). Third, induction of a heat shock inducible *hsp-16p::GFP* reporter, which shares the same *unc-54* 3’UTR and polyA signal as the *gpdh-1p::GFP* reporter, is induced to wild type levels in *cpf-2(dr99)* and *symk-1(dr88)* (Fig. S8). Therefore, inhibition of *cpf-2* and *symk-1* does not generally block mRNA cleavage and polyadenylation.

To test if the Nio mutants alter transcriptional termination and/or recognition of specific polyA signal sequences, we asked if they could suppress phenotypes associated with either *lin-15(n765ts)* or *kin-20(ox423)*. Previous studies demonstrated that inhibition of some core components of the 3’ mRNA cleavage complex suppresses these alleles specifically due to alterations in the ability of the complex to appropriately recognize DNA encoded alternative polyadenylation signal sequences (CUI *et al*. 2008; LABELLA *et al*. 2020). We found that the Nio allele of *cpf-2* completely suppressed the multivulval phenotype *of lin-15(n765)* (Fig. 5). Additionally, *symk-1* and *cpf-2* Nio alleles both suppressed the motility defects associated with the *kin-20* LOF mutation as well or better than the previously identified *kin-20* suppressor *cpsf-4(ox645)* (Fig. 5). Together, these data show that *cpf-2* and *symk-1* Nio alleles both act to inhibit 3’ mRNA cleavage and can disrupt alternative polyadenylation for these specific transcripts.

**Figure 5.**
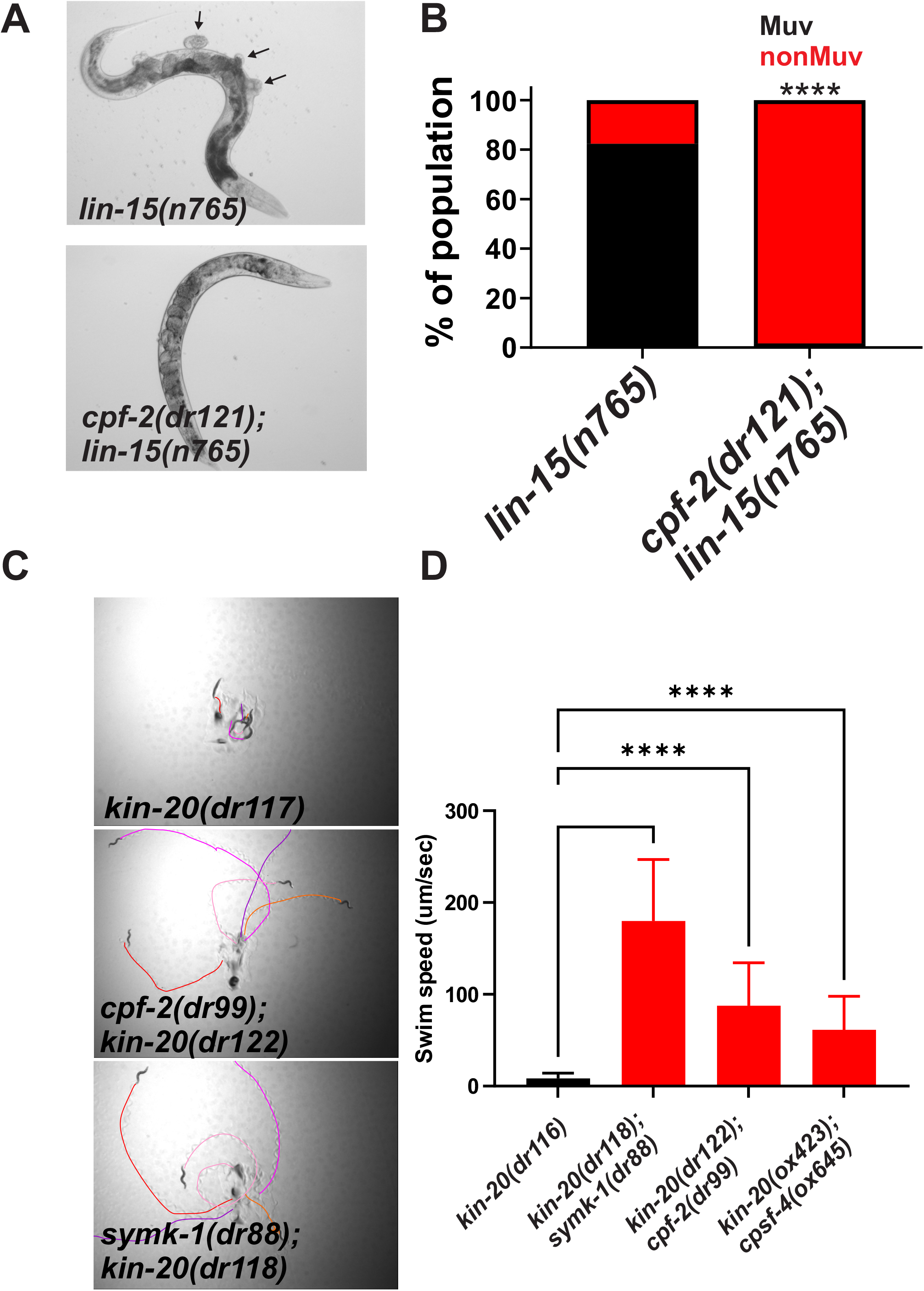
*cpf-2* and *symk-1* suppress mutant phenotypes that depend on alternative polyadenylation. A) Transmitted light images of *lin-15* or *lin-15; cpf-2* double mutants. Arrows in A point to multiple ectopic vulva which produces the multivulval or Muv phenotype. B) Quantification of the % of Muv and nonMuv animals from the indicated genotypes. ****-p<0.0001, Fishers exact test. N=102 animals for *lin-15* and 120 animals for *cpf-2;lin-15*. C) Movement tracks for day 1 adult animals of the indicated genotypes. Each color represents and individual track. D) Swim speed for day 1 adult animals of the indicated genotypes. Data shown are mean ± S.D for 10-24 animals per genotype. ****-p<0.0001, One way ANOVA with Dunn’s multiple comparison test.

To further test this hypothesis, we examined if the Nio phenotype was only caused by disruption of *cpf-2* and *symk-1* or if inhibition of other genes in the 3’ mRNA cleavage complex could also cause a Nio phenotype. 16/17 genes in the 3’ mRNA cleavage complex have clear orthologs in *C. elegans (STEBER et al. 2019)*. We obtained RNAi clones for 12/17 genes and performed post-developmental feeding based RNAi in the *gpdh-1p::GFP* reporter strain. Following exposure to hypertonic stress, knockdown of *suf-1, cpsf-2, cpsf-4*, and *rbpl-1*, in addition to *cpf-2* and *symk-1* (6/12 tested), gave rise to a Nio phenotype (Fig. 6). Therefore, the Nio phenotype is not restricted to disruption of *cpf-2* and *symk-1* specifically but is caused by inhibition of multiple genes in the 3’ mRNA cleavage complex. Taken together, these data are consistent with a model in which inhibition of *cpf-2* and *symk-1* prevent hypertonic induction of *gpdh-1* due to the inability to recognize appropriate polyA signal sequences in a gene(s) required for activation of the hypertonic stress response.

**Figure 6.**
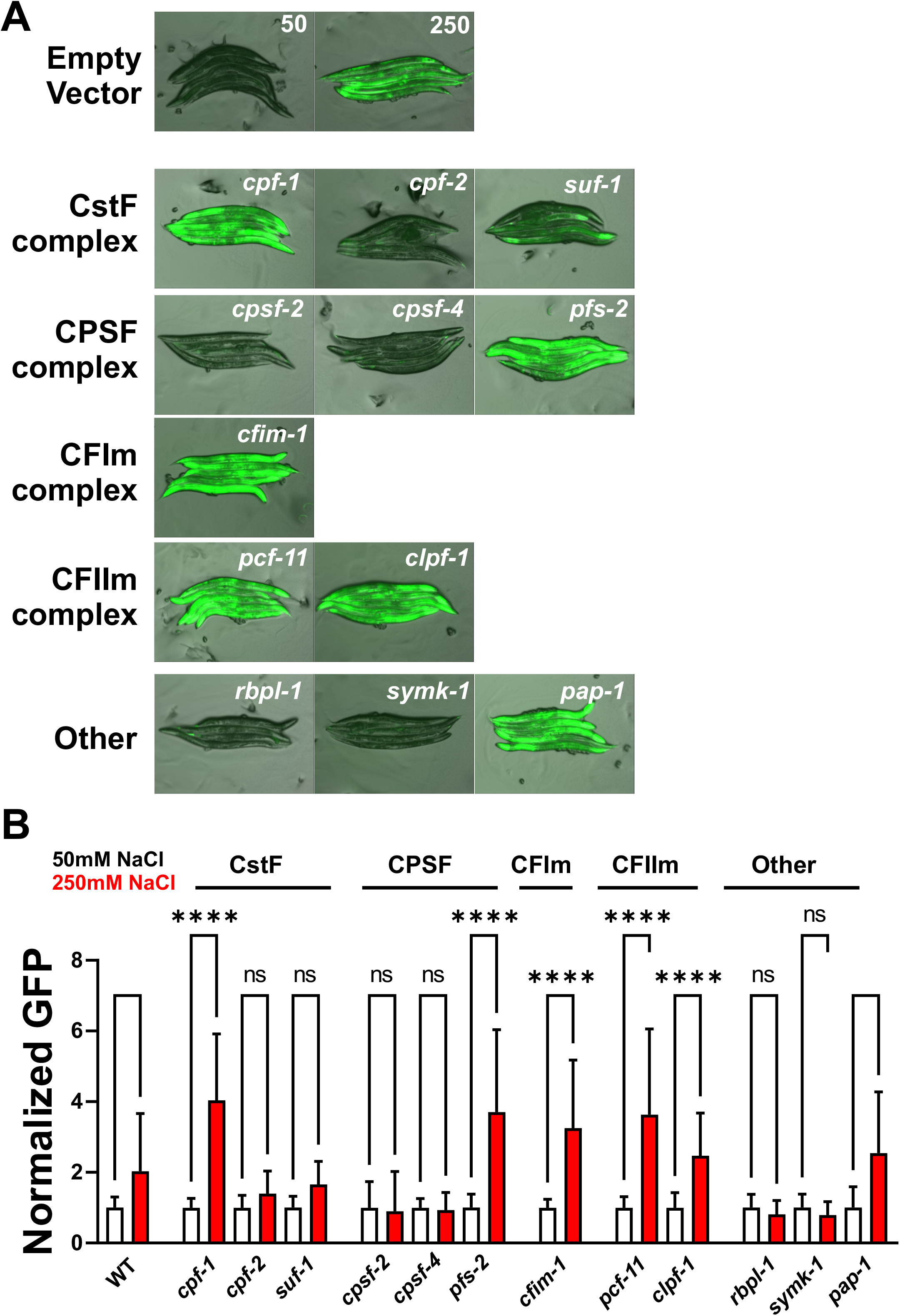
Inhibition of a subset of APA complex genes causes a Nio phenotype. A) Merged GFP and transmitted light images of *drIs4* animals fed either *empty vector(RNAi)* or RNAi against the indicated APA complex genes. Only images for animals exposed to 250 mM NaCl for 24 hours are shown for APA RNAi treatments. B) COPAS Biosort quantification of normalized GFP levels for each APA RNAi treatment following exposure to either 50 mM NaCl or 250 mM NaCl. Data shown are mean ± S.D. N=14-61 animals per sample. ****-p<0.0001, ns-p>0.05, One way ANOVA with Sidak’s multiple comparison test.

## Discussion

Phenotypes associated with loss of components of the 3’ mRNA cleavage complex, particularly at the organismal level, are poorly understood due to the fact that loss of function mutants in most members of this complex are lethal (STEBER *et al*. 2019). One recent exception to this is in *C. elegans*, where unbiased genetic screens can isolate non-null hypomorphic alleles of genes whose null phenotype is lethal. For example, APA of the *unc-44*/ankyrin mRNA utilizes a distal polyA site in both GABA neurons and hypodermal cells to produce ‘Giant’ ankyrin (CHEN *et al*. 2017; LABELLA *et al*. 2020). In neurons, this isoform switch from normal ankyrin to Giant ankyrin at the correct time is critical to restrain juvenile patterns of neuronal development. The protein kinase A KIN-20 allows transcription to bypass the proximal polyA site and produce giant ankyrin. In *kin-20* mutants, giant ankyrin is not produced, GABA neurons develop abnormally, and mutant animals are severely uncoordinated. Hypomorphic alleles of several 3’ mRNA cleavage complex genes were isolated in a genetic screen for mutants that suppress the *kin-20* Unc phenotype. These mutants disfavor the utilization of the proximal polyA site in *unc-44*, leading to the production of giant ankyrin. Likewise, in the hypertonic stress response, APA of a specific target via regulation of the 3’ mRNA processing complex could be required for appropriate execution of this physiological response. In this respect, one possible APA target is *gpdh-1*. However, evidence from thousands of RNAseq datasets does not reveal evidence for APA of *gpdh-1* itself (STEBER *et al*. 2019) and genetic studies show that *gpdh-1* null alleles can still activate hypertonic adaptation (URSO *et al*. 2020). Instead, we hypothesize that 3’ mRNA cleavage performs a similar function in the hypodermis and intestine to that in the neurons, where it regulates polyA site usage of a signaling gene that is an essential signaling component of the hypertonic stress response. Alternatively, the function of the 3’ cleavage complex could regulate isoform switching for a suite of genes, such as extracellular matrix genes, whose collective function appears to be critical for the hypertonic stress response (ROHLFING *et al*. 2010; ROHLFING *et al*. 2011; DODD *et al*. 2018). Arguing against this second possibility is our observation that removal of the SYMK-1 protein in young adults after deposition of the ECM and immediately before exposure to hypertonicity is sufficient to block the hypertonic stress response. Defining the APA targets of 3’ mRNA cleavage complex in the *C. elegans* hypodermis using RNAseq and evaluating their functional roles in the Nio phenotype represents an important future direction and will help to further distinguish between these and other models. Subcellular localization could also play a major role in the regulation of mRNA cleavage. Both CPF-2 and SYMK-1 are expressed in all observable somatic and germline nuclei in *C. elegans*. We found that hypertonic stress causes both proteins to undergo reorganization into subnuclear foci. The function of these foci is unclear.

Recently, hyperosmotic stress was found to result in the phase separation of the APA factor CPSF6 (JALIHAL *et al*. 2020), which leads to functional impairment of mRNA cleavage and polyadenylation. Whether or not this relocalization of the APA complex in *C. elegans* is related to its role in the Nio phenotype is not known. We found that introducing the G42E missense Nio allele into CPF-2 did not impact its ability to form subnuclear foci in response to hypertonicity or to colocalize with SYMK-1. Further studies will be needed to determine if hypertonic stress leads to functional impairment of transcriptional termination in *C. elegans* as it does in mammalian cells and whether or not large-scale termination events or target-specific APA contributes to the Nio phenotype.

In the same genetic screen that identified *cpf-2* and *symk-1*, we also identified null alleles in the O-GlcNAc transferase *ogt-1* (URSO *et al*. 2020). The discovery of all three genes in the same genetic screen suggests they might act together as part of the same signaling complex. However, our data suggest this may not be the case. First, *ogt-1* regulates *gpdh-1* expression via a post-transcriptional mechanism while *cpf-2* and *symk-1* function at the transcriptional level. Second, while both OGT-1 and CPF-2/SYMK-1 are nuclear proteins and form subnuclear foci in response to hypertonicity, these foci are spatially distinct. Despite these differences, there could be regulatory interactions between *ogt-1* and the 3’ cleavage complex. For example, a recent study shows that the TPR domain of mammalian OGT interacts with the 3’ cleavage complex components PCF11 (CFIIm complex) and CPSF1 (STEPHEN *et al*. 2021). Interestingly, it is the TPR domain, and not the catalytic activity, of OGT-1 that is required for the hypertonic stress response in *C. elegans* (URSO *et al*. 2020). Studies examining the role of *ogt-1* in the regulation of APA events will be needed to differentiate between these possibilities.

While the occurrence of APA has been known for several years, the regulatory mechanisms that determine which polyA site in a given gene will be utilized remains mysterious. Genetic approaches in organisms such as *C. elegans*, provide a powerful opportunity to study this highly conserved and essential complex in many dynamic, tissue-specific settings. The tools and insights developed in this study will play a major role in future studies aimed at understanding the physiologically relevant targets of 3’ mRNA cleavage and alternative polyadenylation in the hypertonic stress response and investigating the mechanisms that drive polyA site switching in response to dynamic environmental changes.

## Materials and methods

### *C. elegans* strains and culture

Strains were cultured on standard NGM media with *E*.*coli* OP50 bacteria at 20°C unless otherwise noted. Hypertonicity was induced by increased plate NaCl concentration to either 200 or 600 mM NaCl. For the auxin-induced degradation experiments, we included 0.5-1.0 mM Naphthaleneacetic Acid (K-NAA) in the NGM media prior to pouring plates. A list of strains used in this study can be found in Table 1.

**Table 1.** *C. elegans* strains

| Strain | Genotype |
| --- | --- |
| OG119 | *drls4* (*gpdh-1p::GFP*; *col-12p::dsRed2*) |
| OG975 | *nio-3/cpf-2(dr16)*; *drls4* |
| OG996 | *nio-7/symk-1(dr23)*; *drls4* |
| OG1137 | *symk-1(dr88)*; *drls4* |
| OG1165 | *cpf-2(dr99)*; *drls4* |
| OG1200 | *kin-20(dr116)* |
| OG1202 | *kin-20(dr118)*; *symk-1(dr88)* |
| OG1207 | *cpf-2(dr121)*; *lin-15(n765ts)* |
| OG1209 | *cpf-2(dr99)*; *kin-20(dr122)* |
| OG1224 | *wrdSi23* (*eft-3p::TIR-1:F2A*; mTAG BFP2:AID* NLS: *tbb3*'UTR); *drls4*; *symk-1(dr127)* [*symk-1::AID\*::GFP*] |
| OG1226 | *reSi2* (*col-10p::TIR-1:F2A*; mTAG BFP2:AID* NLS: *tbb3*'UTR); *drls4*; *symk-1* (*dr129*) [AID* GFP CRISPR into *symk-1*] |
| OG1227 | *reSi12* (*ges-1p::TIR-1:F2A*; mTAG BFP2:AID* NLS: *tbb3*'UTR); *drls4*; *symk-1* (*dr130*) [AID* GFP CRISPR into *symk-1*] |
| OG1241 | *cpSi171* [*vha-8p::TIR1::F2A::mTagBFP2::AID\*::NLS::tbb-2* 3'UTR]; *drls4*; *symk-1(dr142)* [AID* mNeonGreen CRISPR into *symk-1*] |
| OG1142 | *cpf-2(dr92)* [TagRFP CRISPR at N-terminus of CPF-2] |
| OG1144 | *ogt-1(dr84)*; *cpf-2(dr92)* |
| OG1194 | *symk-1(dr111)* [GFP CRISPR just before NLS] |
| OG1195 | *cpf-2(dr92)*; *symk-1(dr112)* [GFP CRISPR just before NLS] |
| OG1203 | *cpf-2(dr99)*; *gpls1* |
| OG1204 | *symk-1(dr88)*; *gpls1* |
| OG1205 | *cpf-2(dr92 dr119)*; *symk-1(dr112)* [TagRFP::cpf-2 with G42E and *symk-1::GFP*] |

### Genetic methods

L4 stage *drIs4* animals (P_0_) were mutagenized in 0.6 mM N-ethyl-N-nitrosourea (ENU) diluted in M9 for 4 hours at 20°C and Nio mutants were isolated as previously described (URSO *et al*. 2020). Backcrossing of both *nio-3(dr16)* and *nio-7(dr23)* to the wild type *drIs4* strain showed that both mutants were recessive. Trans-heterozygotes between *dr16* and *dr23* exhibited complementation. SNPs in both *dr16* and *dr23* were identified using whole genome resequencing and a Galaxy workflow (URSO *et al*. 2020). Candidate genes were tested using gene-specific RNAi in the *drIs4* background. RNAi was performed as described previously (URSO *et al*. 2020).

### COPAS Biosort Acquisition and Analysis

Day one adults from a synchronized egg lay or hypochlorite preparation were seeded on 50 or 250 mM NaCl OP50 or the indicated RNAi NGM plates. After 18 hours, the GFP and RFP fluorescence intensity, TOF, and EXT of each animal was acquired with the COPAS Biosort. Events in which the RFP intensity of adult animals (TOF >400) was <20 (dead worms or other objects) were excluded from the analysis. The GFP fluorescence intensity of each animal was normalized to its RFP fluorescence intensity. To determine the fold induction of GFP for each animal, each GFP/RFP was divided by the average GFP/RFP of that strain exposed to 50 mM NaCl.

### CRISPR/Cas9 genomic editing

For CRISPR editing, we injected purified and assembled Cas9/gRNA/tracrRNA complexes with the purified repair template (all from IDT) and the *rol-6* marker plasmid into the germlines of day 1 adult hermaphrodites. Editing protocols primarily followed Ghanta et al with some modifications (GHANTA AND MELLO 2020). We found that the high concentrations of nucleic acids frequently led to precipitation of Cas9 complexes and needle clogging during injection. Reducing the input levels of gRNA and tracer RNA to 20ng/µl and repair templates to 25 ng/µl (oligo repair) or 10 ng/µl (PCR-based repair templates) prevented needle clogging and led to similar numbers of editing events. Overall, 10-20 P0s were injected and 2-3 plates with >10 rollers were selected for screening. Since CRISPR editing appears to occur primarily in the distal germline while transgene formation occurs primarily in the proximal germline, we screened 24 non-Rol F1 animals that were younger than the Rol animals for editing events using edit-specific PCR approaches. We isolated homozygous edit-positive animals from 2-3 independent heterozygous P0s. All edits were verified by DNA sequencing. A list of all guide RNAs, repair template oligos/primers, and genotyping primers can be found in Table 2.

**Table 2.**
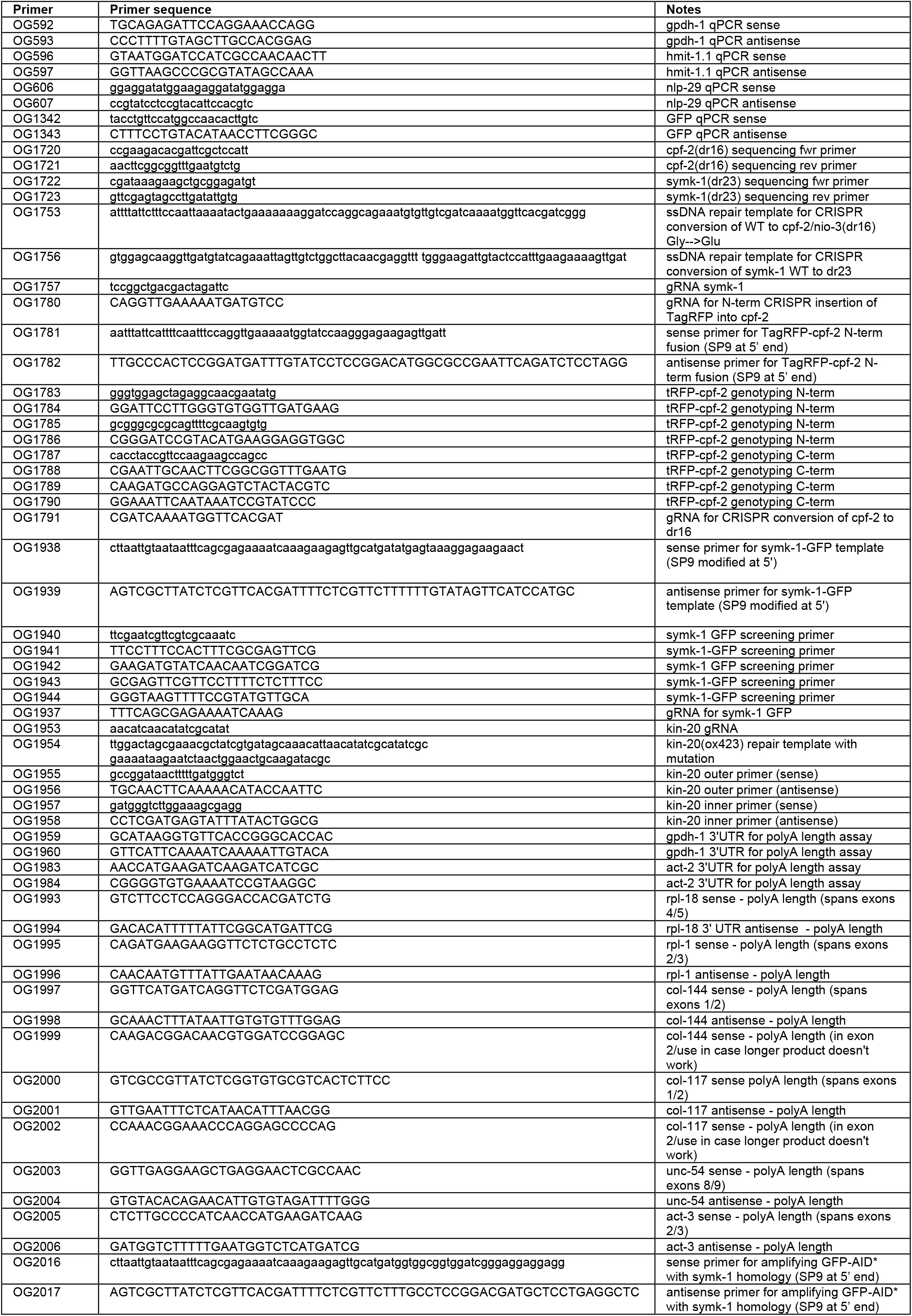
Primers used in this study.

### mRNA isolation, cDNA synthesis, qPCR, and polyA tail analysis

Day one animals were plated on 50 mM or 250 mM NaCl OP50 NGM plates for 24 hours. After 24 hours, 35 animals were picked into 50 µL Trizol for mRNA isolation. RNA isolation followed a combined Trizol/RNeasy column purification method as previously described (ROHLFING *et al*. 2010). cDNA was synthesized from 1 µg total RNA using the SuperScript VILO Master Mix. SYBR Green master mix, 2.5 ng RNA, and the primers listed in Table 2 were used for each qPCR reaction. qPCR reactions were carried out using a BioRad CFX qPCR machine. *act-2* primers were used as a control for all qPCR reactions. At least three biological replicates of each qPCR reaction were carried with three technical replicates per biological replicate. qPCR data was analyzed through ∆∆Ct analysis with all samples normalized to *act-2*. polyA tail length analysis was performed using the Poly(A) Tail-Length Assay Kit (Thermofisher).

### *Symk-1-AID\** alleles and auxin-induced degradation experiments

*symk-1AID*GFP* or *symk-1-AID*mNeonGreen* cassettes were PCR amplified from previously described plasmids (ASHLEY *et al*. 2021) and used as the repair substrate for CRISPR/Cas9 injections into *TIR1; drIs4* strains. TIR1 strains were previously described (ASHLEY *et al*. 2021). AID* alleles of *symk-1* recapitulated the *symk-1* null phenotype (100% emb/larval lethal) when gravid adults expressing TIR1 globally were placed on plates containing either 0.5mM or 1.0mM K-NAA. For tissue-specific Nio experiments, AID* homozygotes eggs were placed onto NGM plates containing 0.5mM or 1mM KAA (Sigma). L4 stage animals were picked to new 1mM auxin plates and aged for 24 hours. Day 1 adults were placed on either 50 mM NaCl NGM / auxin or 250 mM NaCl NGM/auxin plates for 18-24 hours. For temporal experiments, Day 1 adult *symk-1-AID\** were placed on 0.5mM K-NAA for 4 hours prior to transfer. Animals were then imaged and the GFP and RFP signals from the *drIs4* reporter were quantified on a COPAS Biosort.

### Microscopy

For whole worm images, worms were anesthetized (10mM levamisole) and manually arrayed on agar plates for fluorescence microscopy. Images were collected on a Leica MZ16FA stereo dissecting scope with a DFC345 FX camera (Leica Microsystems, Wetzlar, Germany). Images within an experiment were collected using the same exposure and magnification settings.

### *kin-20* and *lin-15* suppression assays

The *ox423* allele of *kin-20* was CRISPR edited into WT, *cpf-2(dr99)*, or *symk-1(dr88)*. Verified *kin-20* homozygotes in each background were placed on a clean NGM plate as a Day 1 adult. Thirty seconds of movement was video captured using a Leica M205 microscope and DFC camera at 15 frames per second. These data were analyzed for multiple kinetic and kinematic parameters using WormLab (MBF Biosciences).

For *lin-15* suppression, *cpf-2(dr99)* and *symk-1(dr88)* heterozygous males were crossed into *lin-15(n765ts)* at the permissive temperature (16ºC) using standard genetic crossing and genotyping methods. For unknown reasons, the *symk-1(dr88)* allele was synthetic lethal with *lin-15(n765ts)*, preventing further analysis. Animals were analyzed for the presence of a multivulval phenotype at the restrictive temperature (20ºC).

## Statistical Analysis

Comparisons of means were analyzed with either a two-tailed Students t-test (2 groups) or ANOVA (3 or more groups) using the Dunnett’s or Tukey’s post-test analysis as indicated in GraphPad Prism 7 (GraphPad Software, Inc., La Jolla, CA). p-values of <0.05 were considered significant.

## Data Availability Statement

Strains and plasmids are available upon request. The authors affirm that all data necessary for confirming the conclusions of the article are present within the article, figures, and tables.

## Acknowledgements

We thank Maria Veroli Noguera, Ph.D. (Pitt) and Robert G. Kalb, M.D. (Northwestern) for critical reading and review of the manuscript prior to submission and Marco Mangone, Ph.D. (Arizona State University) for helpful discussions.

## Supplemental Figure legends

**Figure S1. *cpf-2* and *symk-1* RNAi causes a Nio phenotype**. *dris4* animals were fed *cpf-2* or *symk-1* RNAi from the L1 stage. Day 1 adult animals were exposed to 50 or 250 mM NaCl. GFP induction was quantified on a COPAS Biosort. Data shown are mean ± S.D. N=122-376 animals per sample. ****-p<0.0001, One way ANOVA with Sidak’s multiple comparison test.

**Figure S2. *nio-7(dr23)* and *nio-3(dr16)* Nio phenotypes are rescued by *symk-1* and**

***cpf-2* genomic DNA transgenes**. A) *nio-7(dr23)* animals were injected with 20ng/µl of purified WRM0619bH10 containing the entirety of the *symk-1* genomic locus. Day 1 adult animals were exposed to 50 or 250 mM NaCl. GFP induction was quantified on a COPAS Biosort. Data shown are mean ± S.D. N=25-51 animals per sample. ****- p<0.0001, ns-p>0.05. One way ANOVA with Sidak’s multiple comparison test. B) *nio-3(dr16)* animals were injected with 20ng/µl of a purified 5.1Kb PCR product containing the entirety of the *cpf-2* genomic locus. Day 1 adult animals were exposed to 50 or 250 mM NaCl. GFP induction was quantified on a COPAS Biosort. Data shown are mean ± S.D. N=25-51 animals per sample. ****-p<0.0001, ns-p>0.05. One way ANOVA with Sidak’s multiple comparison test.

**Figure S3 – A *symk-1::GFP::AID\** allele is functional rapidly degraded in the presence of 1mm K-NAA**. GFP images from animals carrying a globally expressed TIR1 transgene and homozygous CRISPR knock-in allele of the GFP::AID* cassette into the C-terminal region of symk-1. Following exposure to 1mM K-NAA, GFP expression is undetectable in all tissues after 3 hours.

**Figures S4. The *symk-1::GFP::AID\** allele is functional**. A) Overlayed GFP and transmitted light images of drIs4 worms that are WT or homozygous for either the *symk-1(dr88)* or *symk-1::GFP::AID\** insertion at the endogenous *symk-1* locus. Animals express TIR1 in the indicated tissues but were not exposed to auxin. B) COPAS Biosort quantification of GFP expression from animals shown in A. Data shown are mean ± S.D. N=71-130 animals per sample. ****-p<0.0001, One-way ANOVA with Dunn’s multiple comparison test.

**Fig S5 - CPF-2 and SYMK-1 hypertonic foci are distinct from OGT-1 foci**. A-C) Hypodermal nuclei (arrows) in animals homozygous for a *TagRFP::cpf-2* (A) and *ogt-1::GFP* alleles (B) form subnuclear foci upon exposure to 600 mM NaCl that fail to co-localize (C). However, TagRFP::cpf-2 (D) and symk-1::GFP (E) exhibit complete colocalization (F).

**Fig S6 - *cpf-2* and *symk-1* Nio mutants do not alter the RNA:DNA ratio**. 10 day 1 adult animals were placed in 12 µl PCR lysis buffer containing 1mg/ml proteinase K, incubated at 60ºC for 1 hour, and then inactivated at 95ºC for 15 minutes. The resulting lysate was used to measure DNA and RNA using the high sensitivity Qubit fluorimetry assay.

**Fig S7 - *cpf-2* and *symk-1* do not alter polyA length/patterns for specific mRNAs**.

A) PCR amplification strategy for measuring polyA mRNA length. PCR reaction 1 shows that the 5’ primer is suitable for amplification. PCR reaction 2 amplifies from the 3’ end of the polyA tail. B)PCR reactions 1 and 2 for eight distinct mRNAs in the indicated genetic background. ‘No RT’ – no reverse transcriptase added to cDNA synthesis reaction.

**Fig S8 - *cpf-2* and *symk-1* do not block *hsp-16p::GFP* induction by heat shock**. A) Images of day 1 adults exposed to either control (20 ºC) or heat shock (3 hours of 35ºC and 18 hours of recovery at 20ºC). B) COPAS Biosort quantification of GFP expression from animals shown in A. Data shown are mean ± S.D, with individual data points shown. N=33-61 animals per sample. ****-p<0.0001, One-way ANOVA with Dunn’s multiple comparison test.

## References

Alt, F. W., A. L. Bothwell, M. Knapp, E. Siden, E. Mather et al., 1980 Synthesis of secreted and membrane-bound immunoglobulin mu heavy chains is directed by mRNAs that differ at their 3’ ends. Cell 20: 293–301.

Ashley, G. E., T. Duong, M. T. Levenson, M. A. Q. Martinez, L. C. Johnson et al., 2021 An expanded auxin-inducible degron toolkit for Caenorhabditis elegans. Genetics 217.

Burkewitz, K., K. P. Choe, E. C. Lee, A. Deonarine and K. Strange, 2012 Characterization of the proteostasis roles of glycerol accumulation, protein degradation and protein synthesis during osmotic stress in C. elegans. PLoS One 7: e34153.

Calfon, M., H. Zeng, F. Urano, J. H. Till, S. R. Hubbard et al., 2002 IRE1 couples endoplasmic reticulum load to secretory capacity by processing the XBP-1 mRNA. Nature 415: 92–96.

Chang, J. W., W. Zhang, H. S. Yeh, M. Park, C. Yao et al., 2018 An integrative model for alternative polyadenylation, IntMAP, delineates mTOR-modulated endoplasmic reticulum stress response. Nucleic Acids Res 46: 5996–6008.

Chen, F., A. D. Chisholm and Y. Jin, 2017 Tissue-specific regulation of alternative polyadenylation represses expression of a neuronal ankyrin isoform in C. elegans epidermal development. Development 144: 698–707.

Clerici, M., M. Faini, R. Aebersold and M. Jinek, 2017 Structural insights into the assembly and polyA signal recognition mechanism of the human CPSF complex. Elife 6.

Clerici, M., M. Faini, L. M. Muckenfuss, R. Aebersold and M. Jinek, 2018 Structural basis of AAUAAA polyadenylation signal recognition by the human CPSF complex. Nat Struct Mol Biol 25: 135–138.

Cui, M., M. A. Allen, A. Larsen, M. Macmorris, M. Han et al., 2008 Genes involved in pre-mRNA 3’-end formation and transcription termination revealed by a lin-15 operon Muv suppressor screen. Proc Natl Acad Sci U S A 105: 16665–16670.

Dodd, W., L. Tang, J. C. Lone, K. Wimberly, C. W. Wu et al., 2018 A Damage Sensor Associated with the Cuticle Coordinates Three Core Environmental Stress Responses in Caenorhabditis elegans. Genetics 208: 1467–1482.

Ghanta, K. S., and C. C. Mello, 2020 Melting dsDNA Donor Molecules Greatly Improves Precision Genome Editing in Caenorhabditis elegans. Genetics 216: 643–650.

Ha, K. C. H., B. J. Blencowe and Q. Morris, 2018 QAPA: a new method for the systematic analysis of alternative polyadenylation from RNA-seq data. Genome Biol 19: 45.

Jalihal, A. P., S. Pitchiaya, L. Xiao, P. Bawa, X. Jiang et al., 2020 Multivalent Proteins Rapidly and Reversibly Phase-Separate upon Osmotic Cell Volume Change. Mol Cell 79: 978–990 e975.

LaBella, M. L., E. J. Hujber, K. A. Moore, R. L. Rawson, S. A. Merrill et al., 2020 Casein Kinase 1delta Stabilizes Mature Axons by Inhibiting Transcription Termination of Ankyrin. Dev Cell 52: 88–103 e106.

Lamitina, S. T., R. Morrison, G. W. Moeckel and K. Strange, 2004 Adaptation of the nematode Caenorhabditis elegans to extreme osmotic stress. Am J Physiol Cell Physiol 286: C785–791.

Lamitina, T., C. G. Huang and K. Strange, 2006 Genome-wide RNAi screening identifies protein damage as a regulator of osmoprotective gene expression. Proc Natl Acad Sci U S A 103: 12173–12178.

Mayr, C., and D. P. Bartel, 2009 Widespread shortening of 3’UTRs by alternative cleavage and polyadenylation activates oncogenes in cancer cells. Cell 138: 673–684.

Pancevac, C., D. C. Goldstone, A. Ramos and I. A. Taylor, 2010 Structure of the Rna15 RRM-RNA complex reveals the molecular basis of GU specificity in transcriptional 3’-end processing factors. Nucleic Acids Res 38: 3119–3132.

Pederiva, C., S. Bohm, A. Julner and M. Farnebo, 2016 Splicing controls the ubiquitin response during DNA double-strand break repair. Cell Death Differ 23: 1648–1657.

Pujol, N., S. Cypowyj, K. Ziegler, A. Millet, A. Astrain et al., 2008 Distinct innate immune responses to infection and wounding in the C. elegans epidermis. Curr Biol 18: 481–489.

Rohlfing, A. K., Y. Miteva, S. Hannenhalli and T. Lamitina, 2010 Genetic and physiological activation of osmosensitive gene expression mimics transcriptional signatures of pathogen infection in C. elegans. PLoS One 5: e9010.

Rohlfing, A. K., Y. Miteva, L. Moronetti, L. He and T. Lamitina, 2011 The Caenorhabditis elegans mucin-like protein OSM-8 negatively regulates osmosensitive physiology via the transmembrane protein PTR-23. PLoS Genet 7: e1001267.

Ruepp, M. D., C. Schweingruber, N. Kleinschmidt and D. Schumperli, 2011 Interactions of CstF-64, CstF-77, and symplekin: implications on localisation and function. Mol Biol Cell 22: 91–104.

Shanbhag, N. M., I. U. Rafalska-Metcalf, C. Balane-Bolivar, S. M. Janicki and R. A. Greenberg, 2010 ATM-dependent chromatin changes silence transcription in cis to DNA double-strand breaks. Cell 141: 970–981.

Shi, Y., D. C. Di Giammartino, D. Taylor, A. Sarkeshik, W. J. Rice et al., 2009 Molecular architecture of the human pre-mRNA 3’ processing complex. Mol Cell 33: 365–376.

Steber, H. S., C. Gallante, S. O’Brien, P. L. Chiu and M. Mangone, 2019 The C. elegans 3’ UTRome v2 resource for studying mRNA cleavage and polyadenylation, 3’-UTR biology, and miRNA targeting. Genome Res 29: 2104–2116.

Stephen, H. M., J. L. Praissman and L. Wells, 2021 Generation of an Interactome for the Tetratricopeptide Repeat Domain of O-GlcNAc Transferase Indicates a Role for the Enzyme in Intellectual Disability. J Proteome Res 20: 1229–1242.

Subramanian, A., M. Hall, H. Hou, M. Mufteev, B. Yu et al., 2021 Alternative polyadenylation is a determinant of oncogenic Ras function. Sci Adv 7: eabh0562.

Urso, S., M. Comly, J. A. Hanover and T. Lamitina, 2020 The O-GlcNAc transferase OGT is a conserved and essential regulator of the cellular and organismal response to hypertonic stress. BioRxiv.

Wang, R., R. Nambiar, D. Zheng and B. Tian, 2018 PolyA_DB 3 catalogs cleavage and polyadenylation sites identified by deep sequencing in multiple genomes. Nucleic Acids Res 46: D315–D319.

Wheeler, J. M., and J. H. Thomas, 2006 Identification of a novel gene family involved in osmotic stress response in Caenorhabditis elegans. Genetics 174: 1327–1336.

Wimberly, K., and K. P. Choe, 2022 An extracellular matrix damage sensor signals through membrane-associated kinase DRL-1 to mediate cytoprotective responses in Caenorhabditis elegans. Genetics 220.

Yuan, F., W. Hankey, E. J. Wagner, W. Li and Q. Wang, 2021 Alternative polyadenylation of mRNA and its role in cancer. Genes Dis 8: 61–72.

Zheng, D., R. Wang, Q. Ding, T. Wang, B. Xie et al., 2018 Cellular stress alters 3’UTR landscape through alternative polyadenylation and isoform-specific degradation. Nat Commun 9: 2268.

