## Supplementary figures and images for "Regulation of the hypertonic stress response by the 3’ mRNA cleavage and polyadenylation complex"

### Supplemental Figure 1

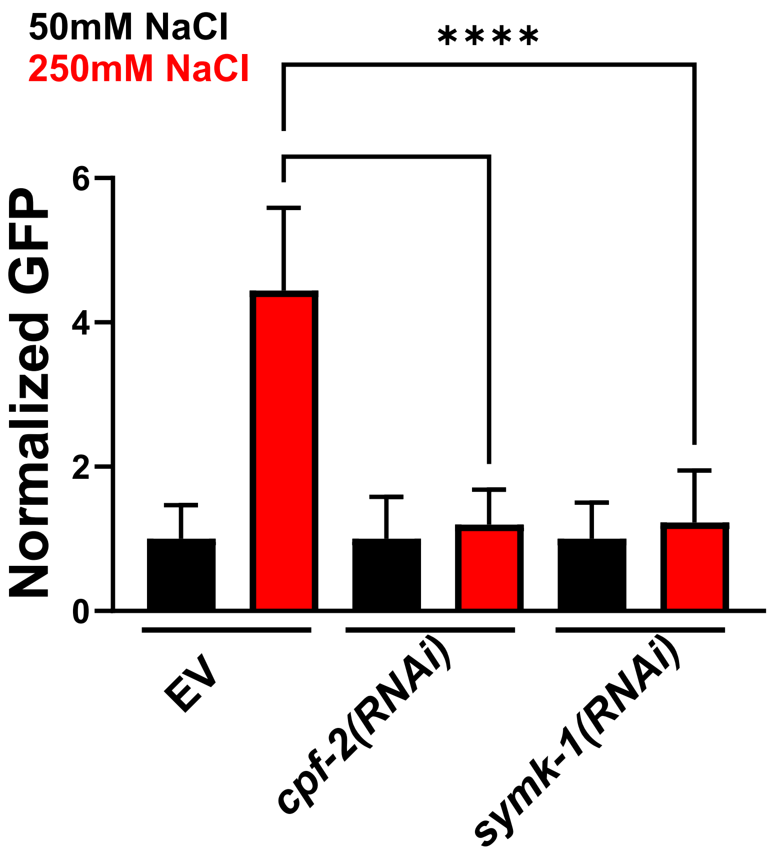

### Supplemental Figure 2

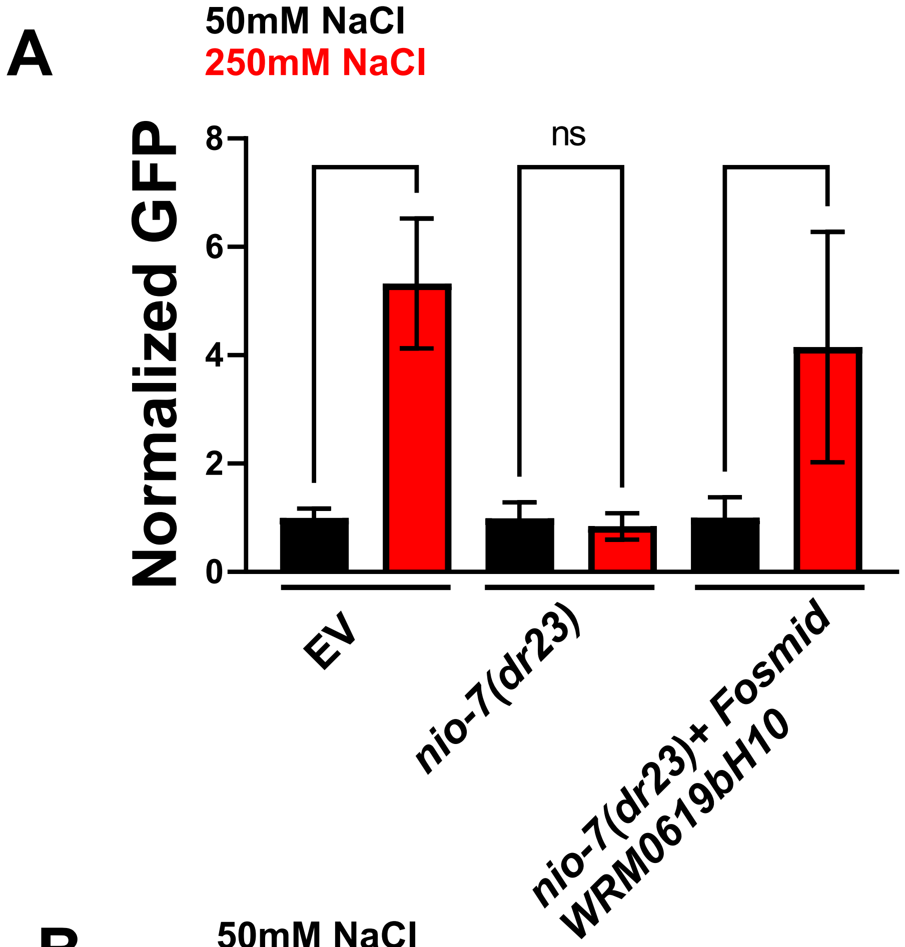

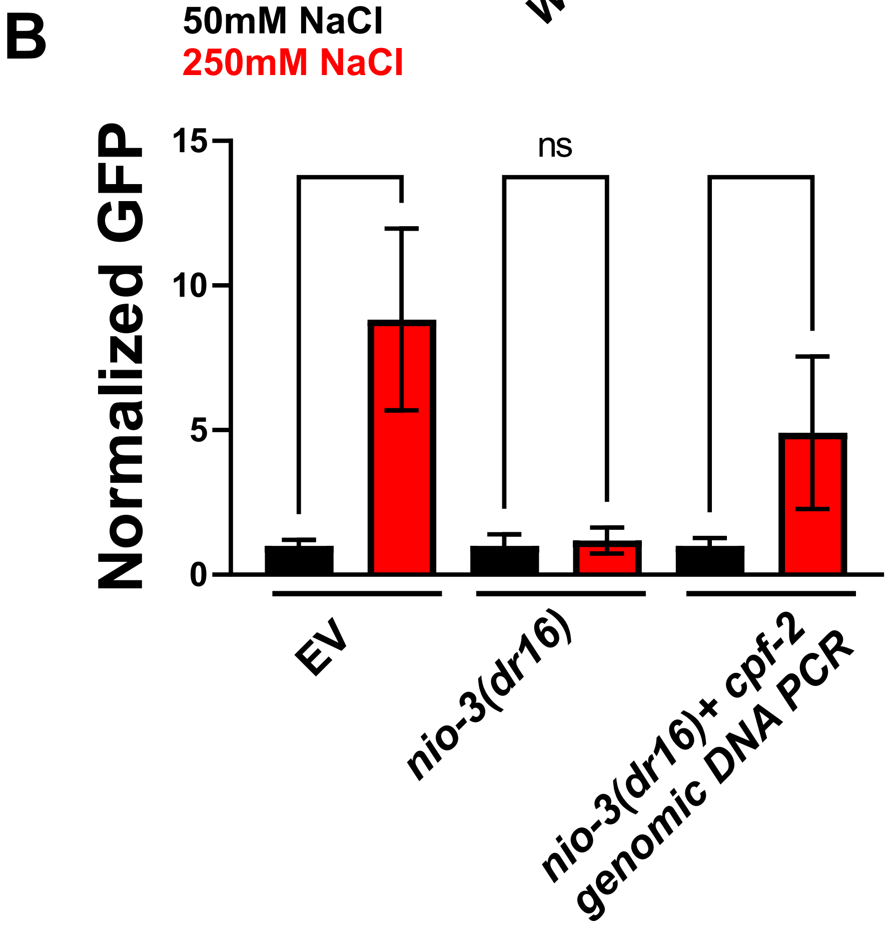

### Supplemental Figure 3

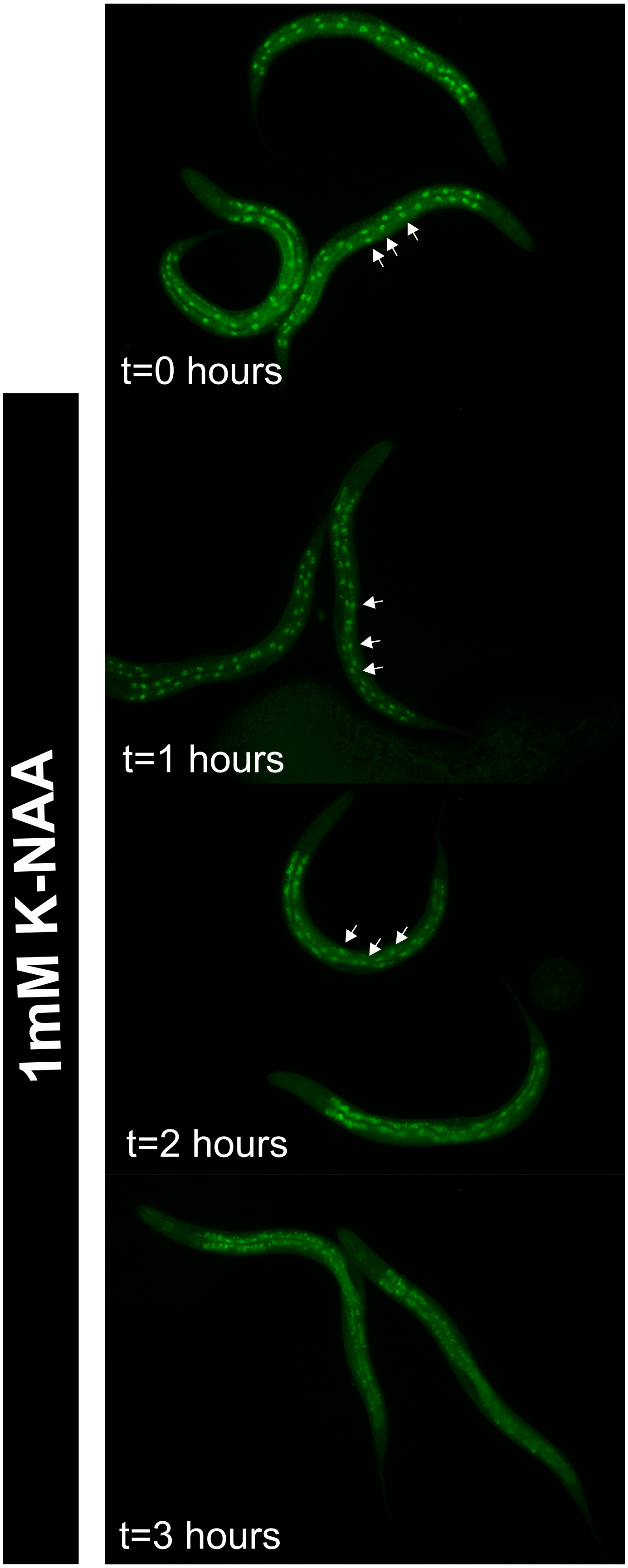

### Supplemental Figure 4

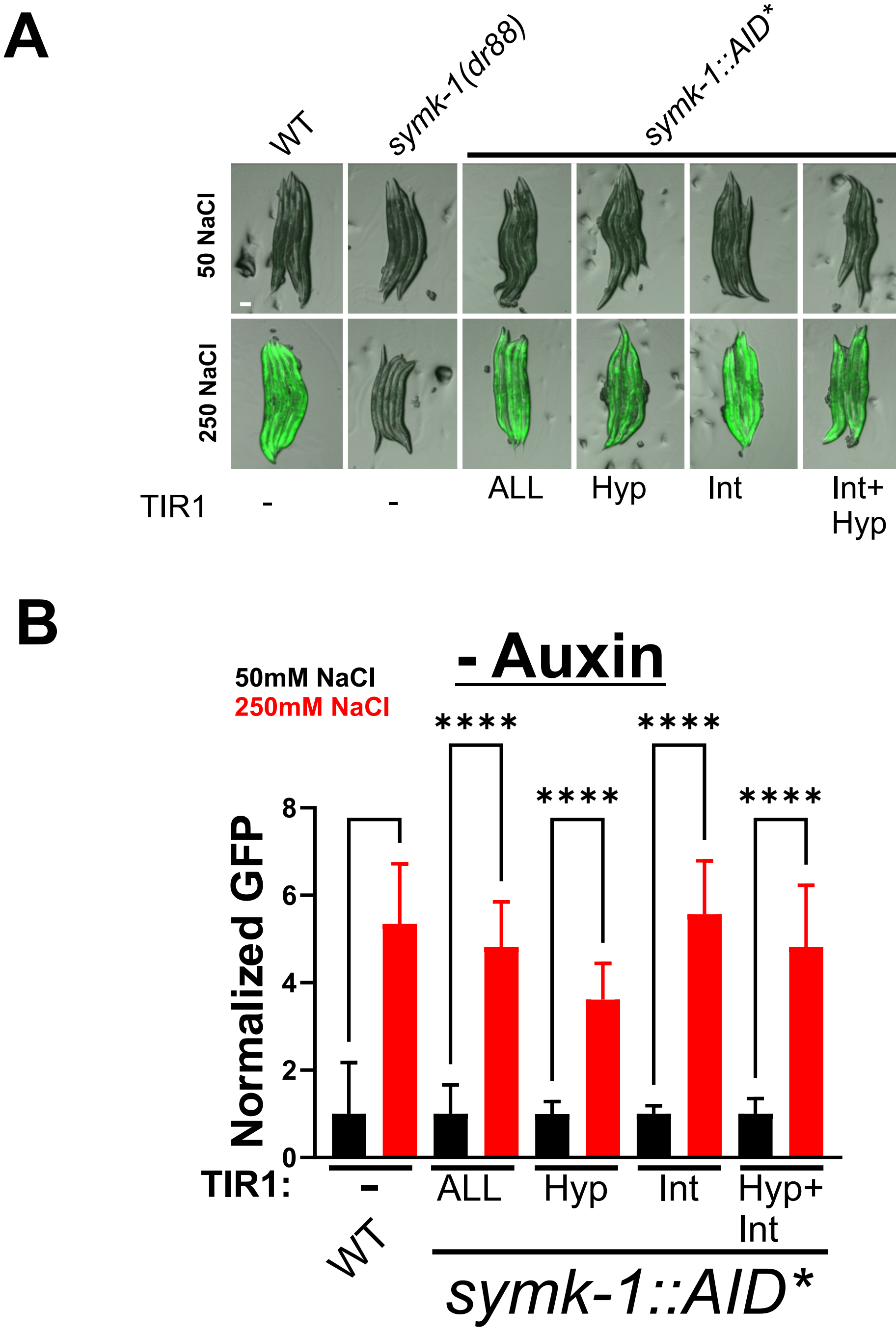

### Supplemental Figure 5

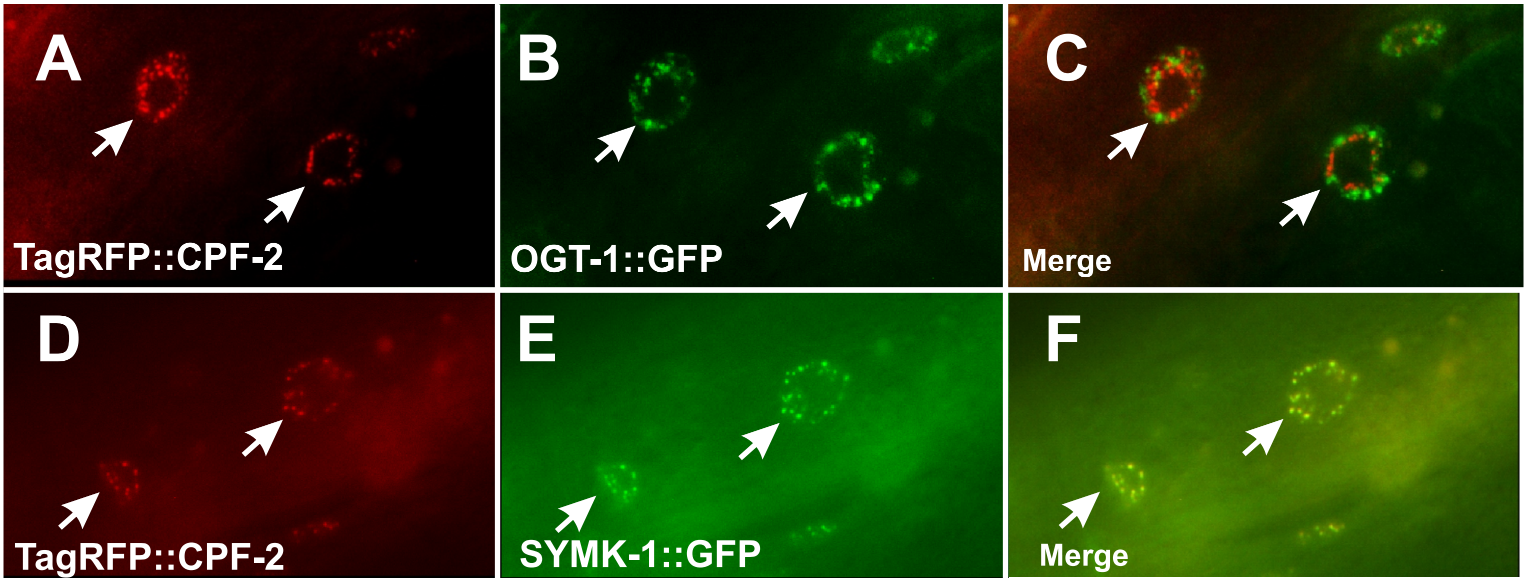

### Supplemental Figure 6

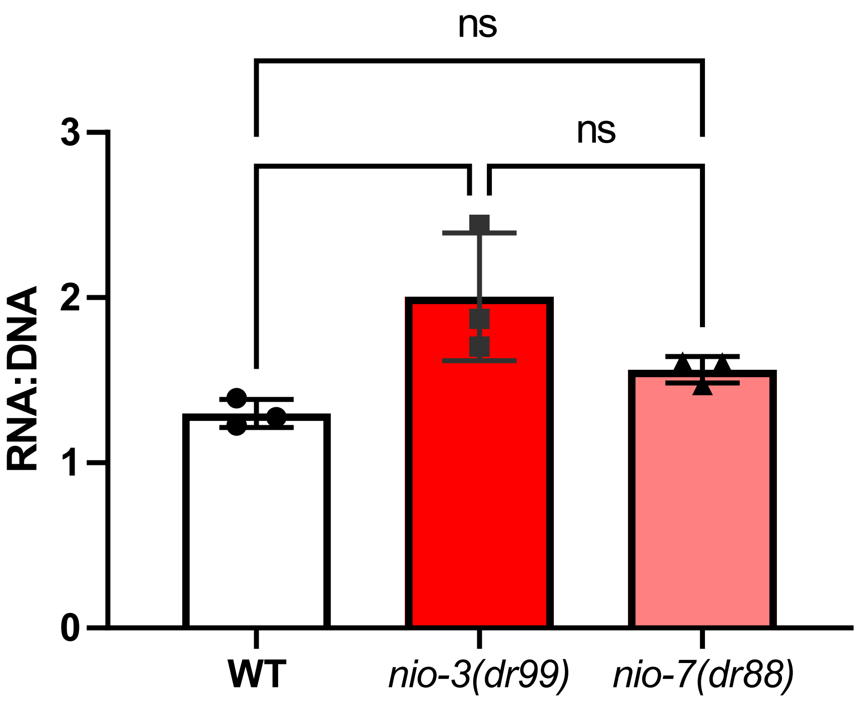

### Supplemental Figure 7

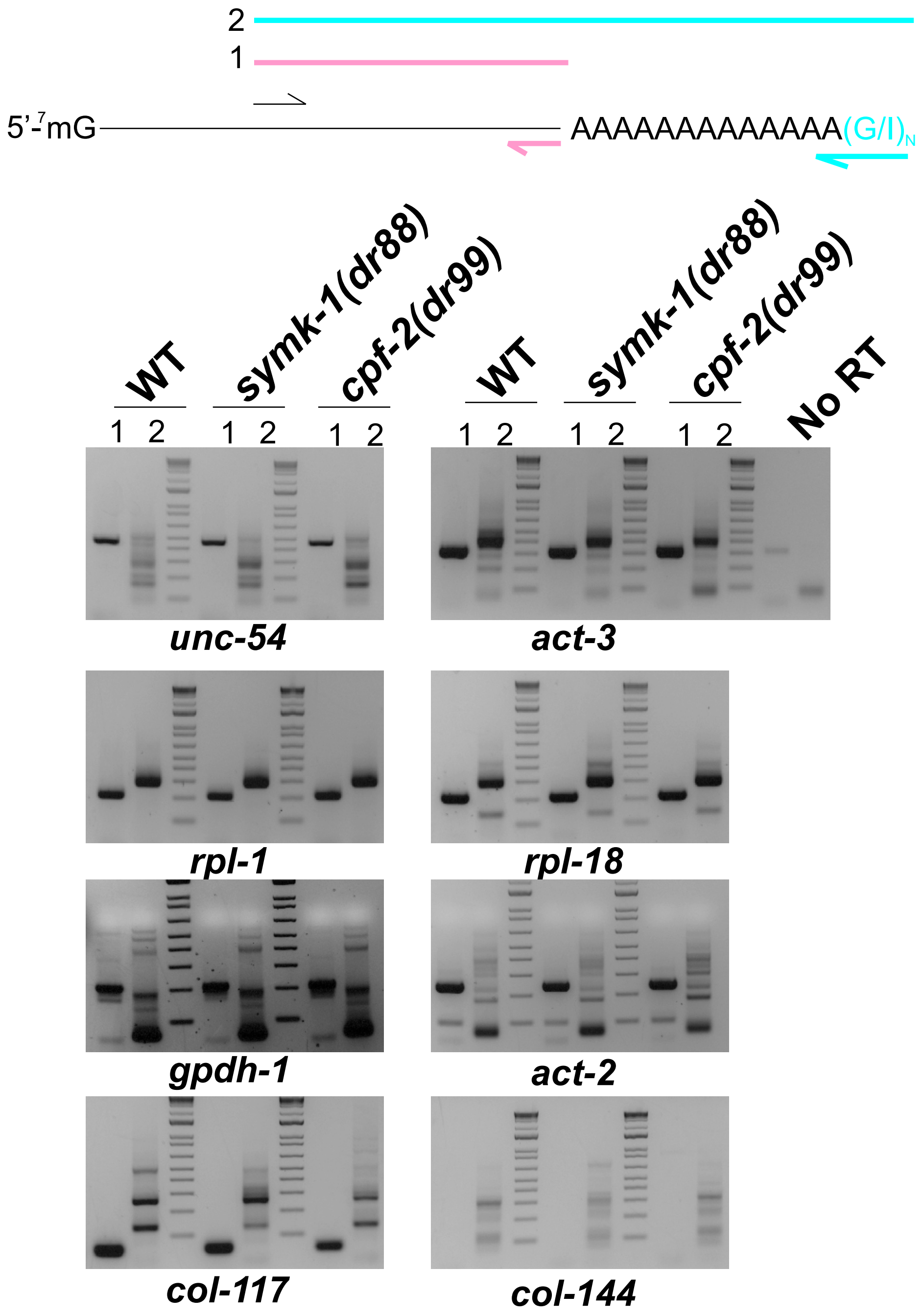

### Supplemental Figure 8

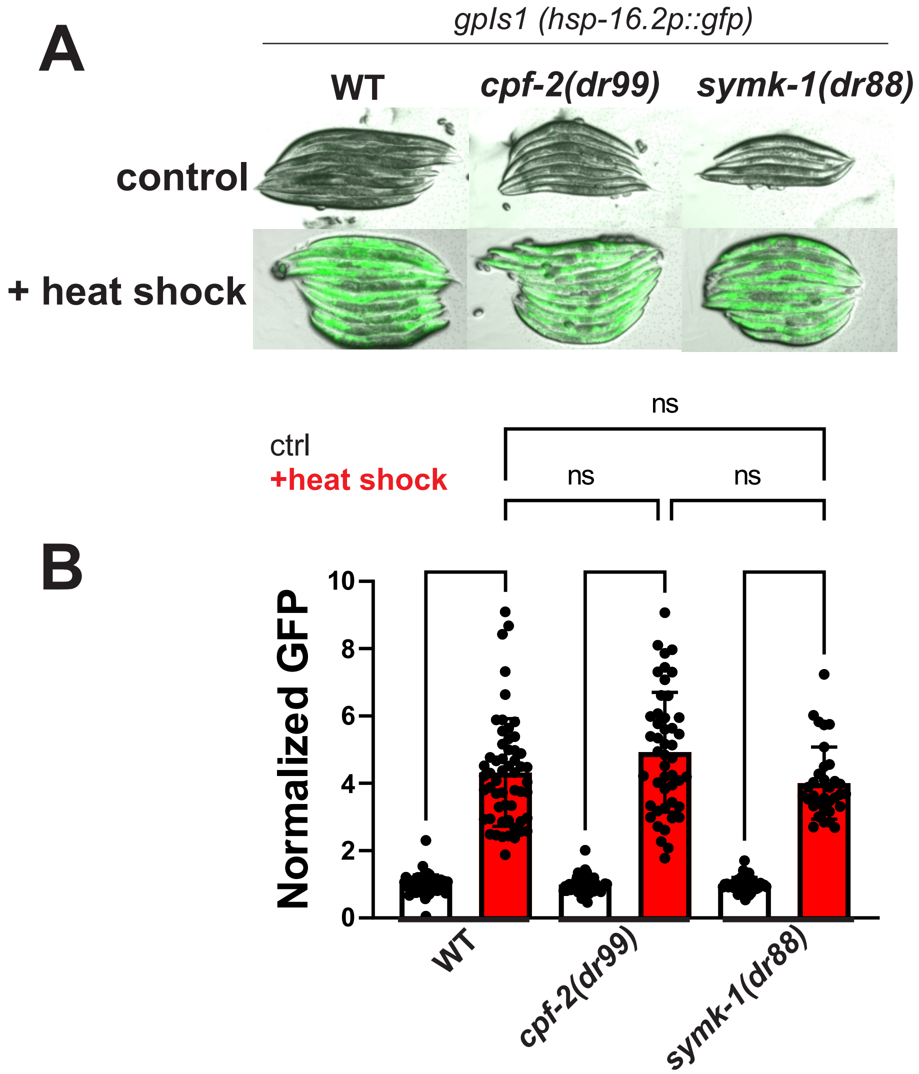
